# Accelerating whole-genome alignment in the age of complete genome assemblies

**DOI:** 10.1101/2024.11.25.625328

**Authors:** Ghanshyam Chandra, Md. Vasimuddin, Sanchit Misra, Chirag Jain

**Author notes:** Emails: { }. Emails: {, }.

## Abstract

Recent advancements in long-read sequencing and assembly methods have ushered in an era of high-quality genome assemblies. Modern assemblies commonly feature megabase-long sequences frequently spanning entire chromosomes. The increase in the assembly contiguity and the reduced number of assembly contigs also implies that whole-genome alignment is no longer an embarrassingly parallel problem. The conventional method of aligning sequences of the query genome in parallel is to utilize a single thread per sequence. This results in poor CPU utilization and long runtimes. In this work, we designed optimizations to accelerate whole-genome alignment on multi-core processors and implemented them in a commonly used aligner, minimap2. Our improvements include a fine-grained parallel chaining method and a fast mechanism for differentiating primary and secondary chains. Our approach accelerates alignment of human, plant, and primate genomes by 1.6*×* to 7.2*×* without compromising accuracy.

## 1 Introduction

Improvements in long-read sequencing technologies have enabled routine assembly of high-quality near-complete genome sequences at affordable costs. Several genome assembly projects, including the Vertebrate Genomes Project [34], Darwin Tree of Life project [33], Bat 1K [40], Human Pangenome Reference Consortium [24], and the Earth Biogenome Project [20], have accelerated the rate at which new assemblies are generated and deposited into public databases. Such efforts also underscore the need for fast and automated solutions to facilitate comparison of these sequences at scale and gather new insights into evolution and inter- and intra-species variation.

Whole-genome alignment has been a fundamental problem in bioinformatics for over two decades, ever since the initial reconstruction of the first animal and plant genomes [3, 4, 28]. Commonly used whole-genome alignment tools feature fast algorithms to compute local alignments while accounting for the presence of complex rearrangements, including inversions, transpositions, and duplications [2]. The classical dynamic programming algorithm requires *O*(*mn*) time, where *m* and *n* are the lengths of two sequences [38]. As *m* and *n* approach the lengths of entire genomes, the exact algorithm becomes too slow in practice. Accordingly, modern whole-genome alignment tools use different heuristic techniques [13, 18, 21, 25, 39]. These heuristic techniques are effective in practice; for example, a systematic benchmarking effort of whole-genome aligners on simulated data showed that the available methods exhibit good accuracy when aligning genomes over close evolutionary distances [8].

Over the last few years, the contig lengths of genome assemblies have dramatically increased. Complete or near-complete assemblies of large genomes are now common [6, 24, 27, 30, 31, 32, 41, 42]. These advancements motivate the need for whole-genome aligners that scale to large, repetitive genomes. Recent efforts on improving the speed of whole-genome alignment involve optimization of data structures [22, 28] and the use of GPU hardware accelerators [10, 11, 12].

Modern processors feature a large number of cores to enable parallel processing. An efficient use of these cores for parallel processing makes a significant difference to the overall runtime. The input to a wholegenome alignment program is a query genome and a reference genome. Each genome can be assumed to be a collection of DNA sequences, e.g., contigs or scaffolds, with non-uniform sequence lengths. The common way of implementing multi-threading in alignment tools, e.g., in minimap2 [21, 22], Mummer4 [28], is to distribute the query sequences across the CPU threads. In this approach, each thread aligns the query sequences assigned to it to the entire reference genome. This approach is simple and efficient for aligning a fragmented genome assembly comprising thousands of contigs, or for aligning a large set of reads. However, the efficiency of this approach drops if the query genome contains a small number of long sequences. For instance, the telomere-to-telomere (T2T) complete genomes contain one sequence per chromosome [32, 42]. As a result, the current approach results in poor work balance among threads. While using minimap2 to align high-quality genome assemblies of various species, we find an average CPU utilization of less than 7% on a 48-core processor, indicating a wide gap between the actual performance and the theoretical peak performance.

In this work, we propose optimizations to accelerate whole-genome aligners on multi-core processors. Our optimizations are applicable to any genome alignment tools which follow *seed-chain-extend* heuristic method [4, 17, 21, 28]. Seed-chain-extend is a widely used heuristic method that involves (i) computing short exact matches between two genomes called *anchors*, (ii) combining groups of anchors into longer, high-scoring *chains* based on their proximity and relative ordering to one another [1, 15], (iii) classifying a chain as either *primary* or *secondary* when the same region of the query genome aligns to two regions in the reference [37], and (iv) computing base-to-base alignments by extending the anchors of each chain. See Figure 1 for an illustration. We implemented our optimizations by modifying the minimap2 source code [21]. We chose minimap2 for implementing and testing the optimizations because (a) minimap2 is among the most widely used whole-genome alignment tools, (b) minimap2 code is well maintained and constantly updated, and (c) it exhibits a good balance between alignment accuracy and computational speed [36].

**Figure 1.**
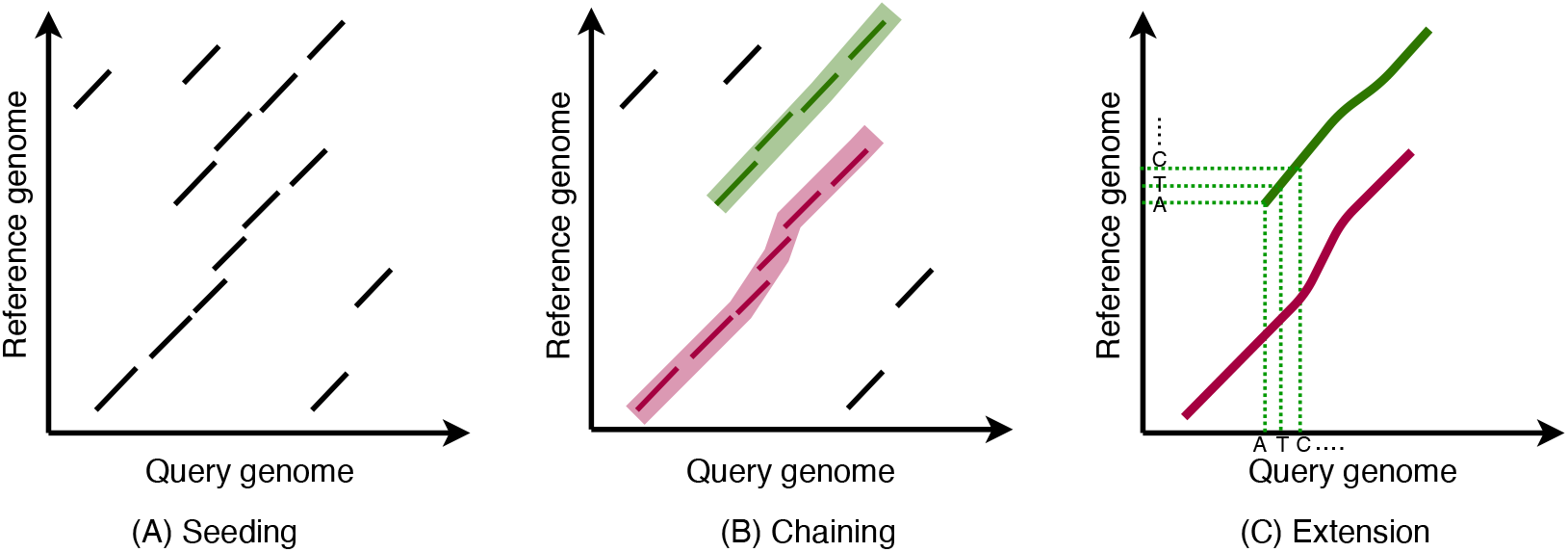
Illustration of the seed-chain-extend heuristic method for computing whole-genome alignment. **(A)** Computation of short exact matches (anchors). Each diagonal line corresponds to an exact substring match between the query and reference genomes. **(B)** Computation of longer chains of multiple anchors. In this example, there are two chains. The two chains have overlapping intervals on the query genome. The chain highlighted in magenta color has a higher score due to more number of anchors and considered as primary, whereas the green chain is considered as secondary. **(C)** Extension of anchors in each chain to obtain base-to-base alignments.

We profiled the percentage contribution of each step in minimap2 to the total runtime of whole-genome alignment. Our results show that all four steps of the seed-chain-extend workflow contribute significantly to the overall runtime. We implemented multiple optimizations to accelerate these steps. First, we developed a fine-grained parallel algorithm for chaining. Our algorithm enables parallel chaining of a single query sequence using multiple threads. Second, we implemented a fast interval tree-based algorithm for classifying chains as primary or secondary. Third, we replaced sequential sorting routines with parallel sorting to accelerate the seeding stage. Fourth, we optimized the extension stage by using the SIMD (single instruction multiple data)-parallel alignment library based on AVX (advanced vector extensions) intrinsics from our previous work [16]. Together, these optimizations result in 1.6 −7.2 *×* speedup for whole-genome alignment while preserving the output accuracy. For example, the original minimap2 implementation requires about 12 hours wall-clock time to align barley genome assembly to the barley reference genome using 48 threads, whereas our implementation, called mm2-plus, takes less than two hours. Accordingly, mm2-plus can offer significant advantage to the users doing genome analysis by reducing genome alignment time and cloud computing costs.

## 2 Results

### Experimental setup

We developed mm2-plus (https://github.com/at-cg/mm2-plus) on top of minimap2 (v2.28) [21]. In order to demonstrate the portability of our improvements, we conducted our experiments on four different CPU architectures: Intel^®^ Xeon^®^ Platinum `8592+` (Emerald Rapids), Intel^®^ Xeon^®^ Platinum `8480L` (Sapphire Rapids), Intel^®^ Xeon^®^ Platinum `6248R` (Cascade Lake), and AMD EPYC™ `7763` (Milan). The hardware specifications of these systems are provided in Supplementary Table S1. We used the following datasets in our evaluation: (a) Barley-Barley, (b) Maize-Maize, (c) Human-Bonobo, and (d) Human-Human. For instance, the Barley-Barley dataset involves aligning a barley genome to another barley genome. Details of these datasets are available in Table 1 and Supplementary Table S2. The contiguity of these genomes is consistent with the quality of the latest genome assemblies, with N50 scaffold lengths exceeding 100 Mbp (Table 1). N50 statistic implies half of the genome bases are represented in sequences of this length or greater. One of these datasets (Human-Bonobo) corresponds to an inter-species genome comparison. The remaining are intra-species genome comparisons. We used the default alignment parameters from minimap2 documentation. The commands are also available in Supplementary Table S3.

**Table 1:**
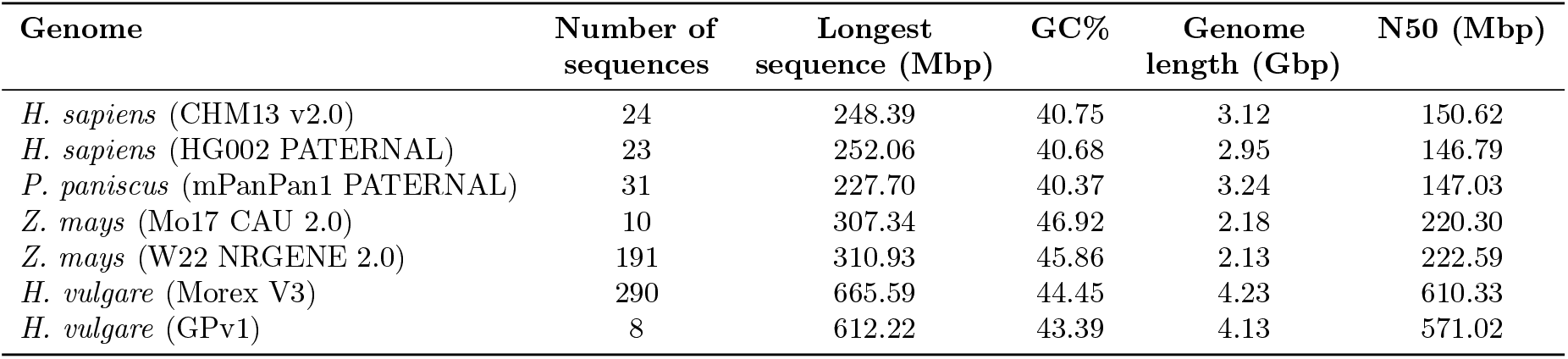
Summary statistics of the genomes used for experiments. We used genomes from *H. sapiens* (human), *P. paniscus* (bonobo), *Z. mays* (maize), and *H. vulgare* (barley) species.

### Runtime profile of minimap2

We profiled the runtime contribution of various steps that are part of the seed-chain-extend workflow of minimap2. We conducted this experiment using three datasets (a) Barley-Barley, (b) Human-Human, and (c) Human-Bonobo. Minimap2 has four key steps: (1) Seeding step for *computing anchors*, (2) Anchor *chaining*, (3) *Marking primary chains*, and (4) Extension step, i.e., *base-to-base alignment*. In the profile, the remaining time apart from these steps is presented as *miscellaneous*. In the step (1), minimap2 identifies common minimizer *k*-mers [35] between the reference and query genomes as anchors. This step is followed by the chaining and the extension steps (Figure 2A). In our experiments done using 48 threads of a Cascade Lake CPU, we profiled runtimes of these steps. The steps consuming more than 5% of the total time in Barley-Barley, Human-Human, and Human-Bonobo genome alignments are shown in Figure 3. The profiling results suggest that the algorithms used in all the four steps must be improved to achieve a significant reduction in the overall runtime. Another important observation is that the CPU utilization ranges from 2.27% to 6.62%, suggesting that most processor cores predominantly remain idle during the program execution due to poor load balance among threads. Total runtime (wall clock time) varies from 11.84 hours (Barley-Barley) to 0.33 hours (Human-Human). Barley genome comparisons require significantly more time because barley genomes are longer and more repetitive compared to human genomes [29].

**Figure 2.**
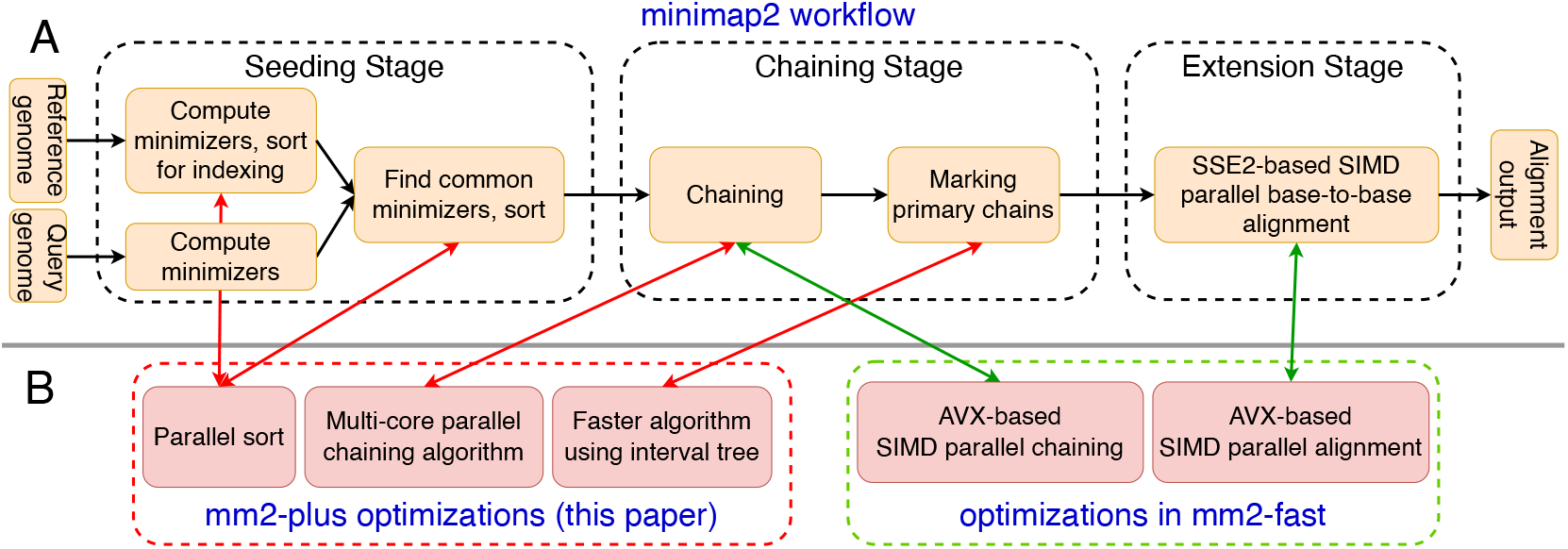
Illustration of the main computational steps in minimap2 and the proposed optimizations. **(A)** The key steps involved in the seed-chain-extend alignment workflow of minimap2. **(B)** The proposed techniques implemented in mm2-plus to accelerate the individual steps. We also used optimizations from our previous work, mm2-fast [16], as shown above.

**Figure 3.**
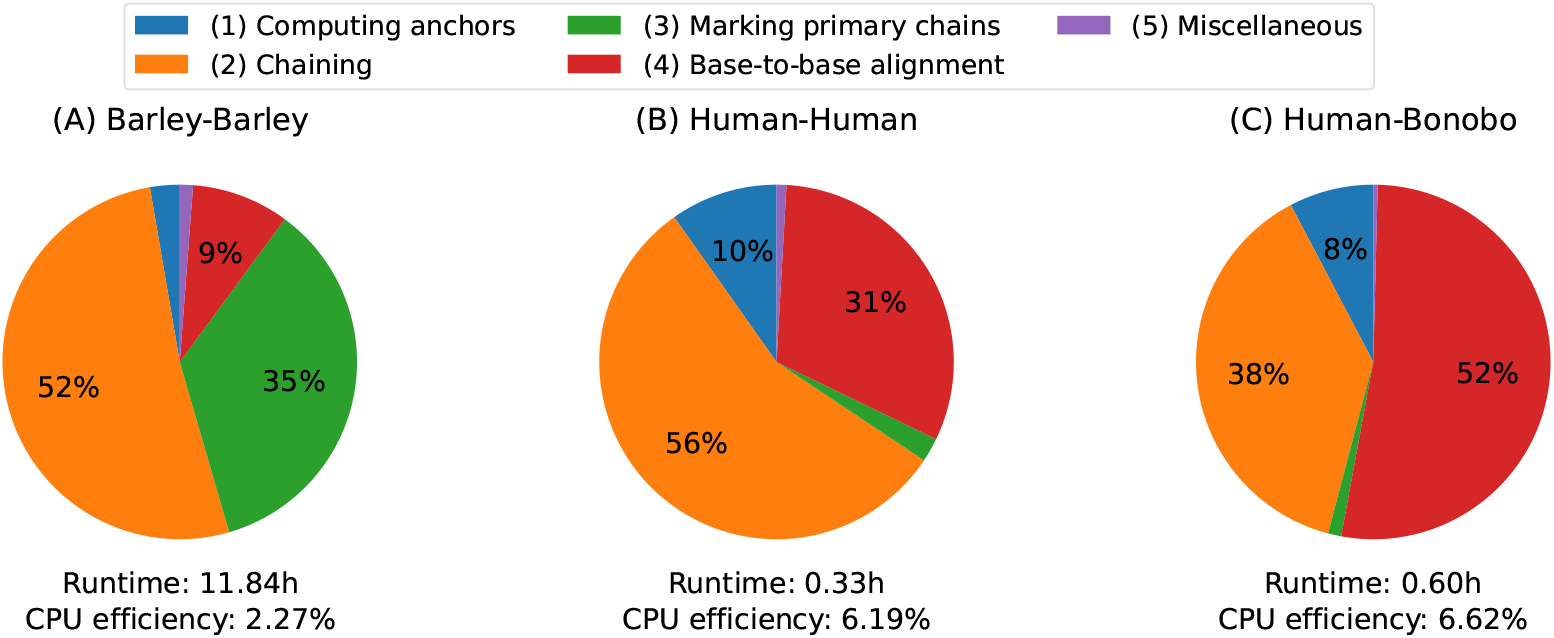
Pie charts to visualize the runtime profiling results of minimap2. We highlight the percentage of runtime spent in various steps. We conducted this experiment using 48 threads. CPU efficiency is calculated as the ratio of actual CPU usage and the maximum possible CPU usage.

### Overview of our optimizations

In the following, we present a summary of the optimizations which we implemented in mm2-plus. More details of these optimizations are provided in Methods.

- *Parallel chaining:* The input to the chaining algorithm is the set of anchors indicating exact matches between the query genome and the reference genome. For a given query sequence, this step is processed sequentially by a single thread in minimap2. We parallelized chaining by partitioning the input set of anchors based on the sequence of the reference genome they lie on. This approach allows independent processing of each partition because a single chain cannot span more than one reference sequence. We use the default chaining algorithm of minimap2 on all partitions, and aggregate the chains identified across all the partitions.
- *Faster algorithm for marking primary chains:* When a segment of a query sequence exhibits multiple alignments to the reference genome, it results in multiple chains overlapping with each other on the coordinates in the query sequence. Minimap2 classifies these chains as either primary or secondary [21]. Among the overlapping chains, the chain with the highest score is marked as the primary chain and the remaining chains are marked secondary chains. The default algorithm in minimap2 to identify primary chains requires quadratic time in the number of chains. This algorithm takes significant time when the number of chains is large, which commonly happens with large plant genomes that have several repeats and mobile elements. To address this limitation, we implemented a faster algorithm that uses sub-quadratic time in practice by utilizing an interval tree data structure.
- *Faster base-to-base alignment using AVX instructions*: Base-to-base alignment also consumes a significant portion of the total runtime because aligning two sequences requires quadratic time in the length of sequences. By default, minimap2 uses SSE2 (128 bit) instructions for parallel processing using SIMD. We leveraged faster alignment code from our previous work [16] that uses AVX2 (256 bit) and AVX512 (512 bit) instructions to increase the parallelism. The AVX instructions are commonly supported in modern AMD and Intel CPU architectures.
- *Parallel sorting* : Sorting is frequently used for various tasks in minimap2, for example, to maintain the ordering of anchors based on their coordinates in the reference genome. Minimap2 uses its own sequential algorithm for sorting. In mm2-plus, we replaced sequencing sorting with a parallel sorting algorithm [5].

We will discuss the runtime advantage of each optimization in the next section. We also checked the accuracy of alignments using variant calling (Methods). Implementing the above optimizations caused a negligible change to the alignment output (Supplementary Table S4). These differences are introduced by subtle changes in the mm2-plus implementation; for example, the sorting algorithm in minimap2 is not stable, whereas the sorting algorithm in mm2-plus is stable.

### Performance evaluation

We compared the runtime of mm2-plus with minimap2 using four datasets: (A) Barley-Barley, (B) Maize-Maize, (C) Human-Human, and (D) Human-Bonobo. We measured the impact of each optimization, labeled as *O*_1_ (faster base-to-base alignment using AVX), *O*_2_ (faster algorithm for marking primary chains), *O*_3_ (parallel chaining), and *O*_4_ (parallel sorting). We compared minimap2 baseline method with different versions of mm2-plus in which we enabled the optimizations progressively. In other words, we compared minimap2, mm2-plus (*O*_1_ enabled), mm2-plus (*O*_1_ + *O*_2_ enabled), mm2-plus (*O*_1_ + *O*_2_ + *O*_3_ enabled), and mm2-plus (*O*_1_ + *O*_2_ + *O*_3_ + *O*_4_ enabled). This experiment is useful to evaluate the advantage of each optimization as well as measure their combined impact. We conducted this experiment using 48 threads on a Cascade Lake CPU. The runtimes are shown in Figure 4.

**Figure 4.**
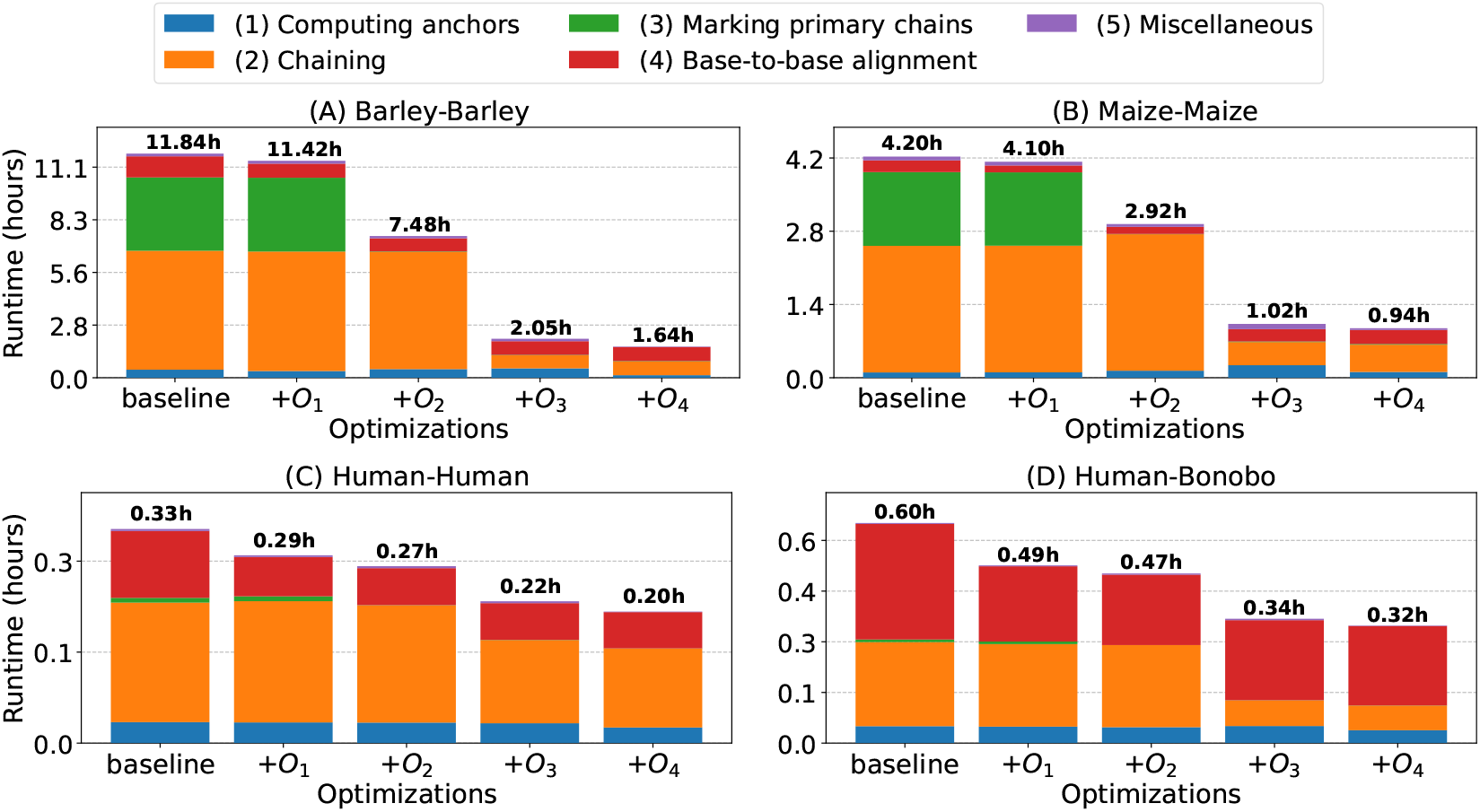
End-to-end runtime (wall-clock time) using minimap2 (baseline) and the different versions of mm2-plus. We progressively enabled the four optimizations *O*_1_, *O*_2_, *O*_3_, and *O*_4_ in mm2-plus for a detailed evaluation. The labeling of the four optimizations is done as *O*_1_ (faster base-to-base alignment using AVX), *O*_2_ (faster algorithm for marking primary chains), *O*_3_ (parallel chaining), and *O*_4_ (parallel sorting). In each bar, we show the runtime contribution of different steps using different colors. We conducted this experiment using 48 threads.

We draw the following conclusions from the results. When all optimizations are enabled, mm2-plus is 1.6 −7.2 *×* faster than minimap2. In particular, we achieved higher speedups using the plant genomes. Our optimizations *O*_2_ and *O*_3_ work better on barley and maize plant genomes because (i) the number of chains computed by minimap2 in these genomes is significantly higher (Supplementary Figure S1) making the interval tree-based implementation for marking primary chains more effective, and (ii) our parallel chaining algorithm based on anchor partitioning works better on these genomes because the anchor positions are scattered across multiple sequences of the corresponding reference genome (Supplementary Figure S2) leading to better thread utilization. Notably, the results highlight that all the optimizations in mm2-plus consistently contribute to reducing runtime across all datasets. The memory-usage of mm2-plus (with all optimizations enabled) and minimap2 are comparable (Supplementary Figure S3).

We also evaluated the performance of mm2-plus when fewer than 48 threads were used (Figure 5). The results indicate that mm2-plus is consistently faster than minimap2, with the highest speedup achieved using ≥ 16 threads. When we use a single thread only, mm2-plus requires much less time for Barley-Barley and Maize-Maize datasets due to the faster method for marking primary chains. As we increase the thread count, we also see the benefit of parallel chaining. Beyond 16 threads, the runtime of mm2-plus nearly saturates across all datasets. The parallel efficiency of mm2-plus chaining algorithm is affected by the number of sequences in the query and reference genomes, and the distributions of partition size when we split the anchors.

**Figure 5.**
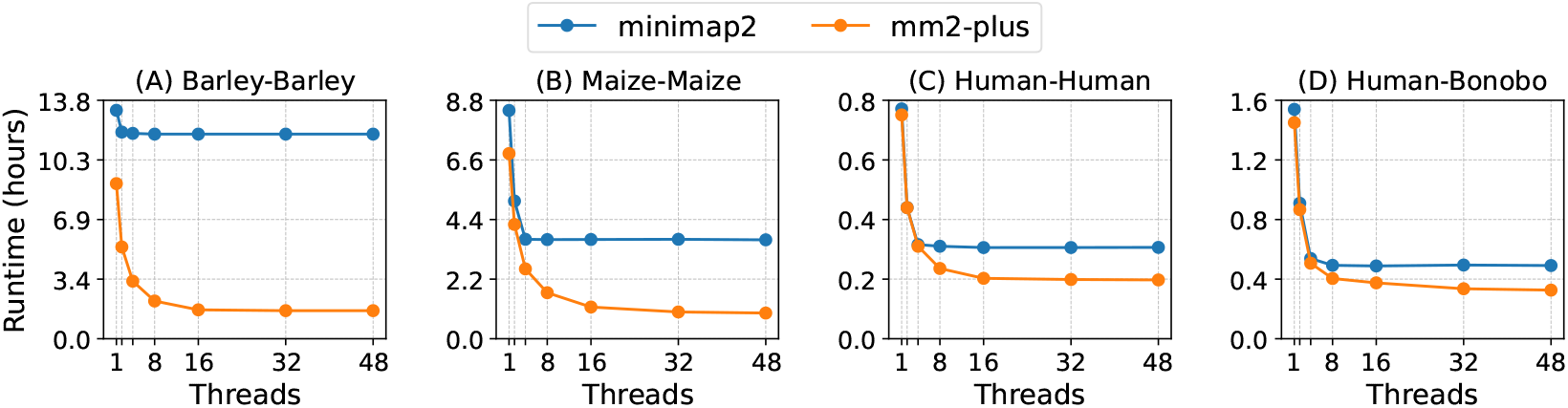
Comparison of mm2-plus and minimap2 runtimes using different number of threads.

### Cross-platform performance and compatibility

Our optimizations in mm2-plus are compatible with a range of Intel and AMD CPUs. To demonstrate this, we benchmarked mm2-plus on four architectures: Intel Emerald Rapids, Intel Sapphire Rapids, Intel Cascade Lake, and AMD Milan. These architectures differ in terms of their core counts, SIMD register widths, CPU frequency, cache size, etc. (Supplementary Table S1). We carried out this experiment using a single socket of each processor. We achieved similar trend of speedups on all architectures (Figure 6), that is, we find a similar trend of higher speedups on plant genomes compared to animal genomes. Our optimizations of parallel chaining, parallel sorting, and the use of faster algorithm for marking primary chains are hardware-agnostic, that is, they generalize to any multi-core processor. The performance of base-to-base alignment using SIMD is affected by the SIMD register width. For example, Intel supports AVX2 (256 bit) and AVX512 (512 bit) instructions, whereas AMD supports only AVX2 instructions. With all the optimizations combined, we achieved higher speedup on Intel processors compared to AMD. On Intel processors, the speedup numbers improved with the newer generations of architecture, that is, from Cascade Lake to Emerald Rapids.

**Figure 6.**
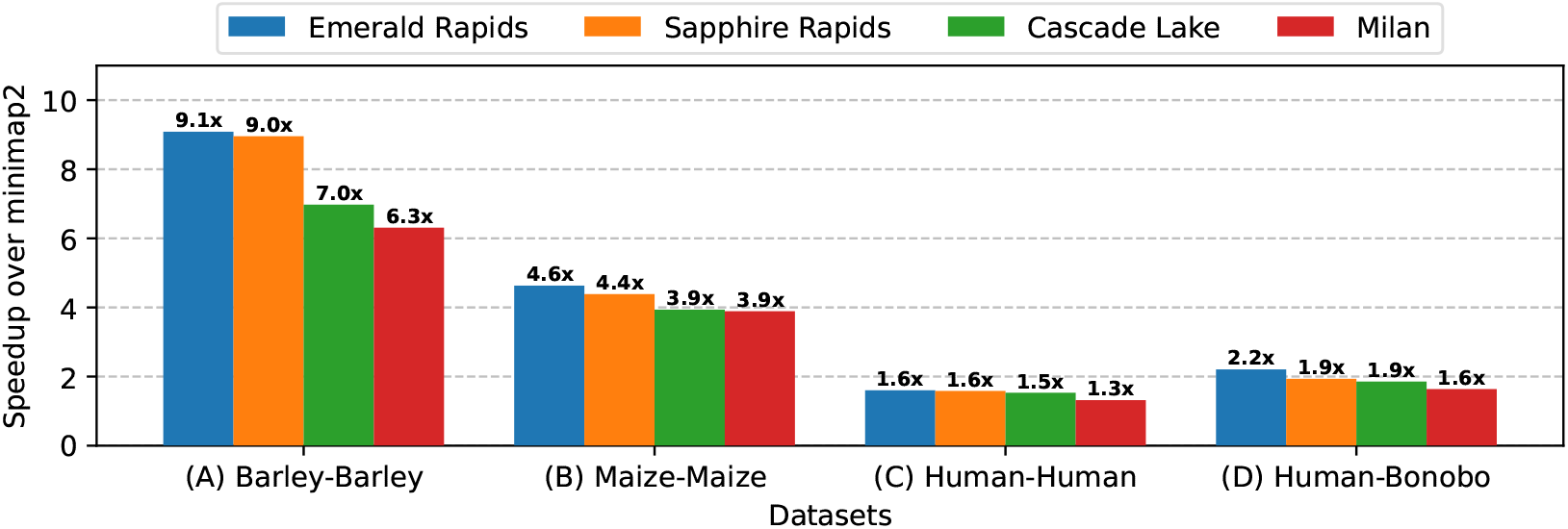
Speedups observed using mm2-plus on different CPU hardware architectures. On each type of hardware, we ran mm2-plus and minimap2 using multiple threads, utilizing all cores of single socket. Emerald Rapids, Sapphire Rapids, and Cascade Lake are different generations of Intel architecture, while Milan is an AMD architecture.

### Support for accelerated execution of other use-cases of minimap2

Minimap2 supports a variety of alignment applications besides whole-genome alignment, including long-read mapping for genome resequencing and all-vs-all read alignments for *de novo* assembly. We have ensured that mm2-plus can be useful for faster execution of these applications too. Accordingly, we ported all optimizations from our earlier work, mm2-fast [16], to mm2-plus as well. These included SIMD-parallel chaining and SIMD-parallel base-to-base alignment [16]. We further added dynamic load balancing to account for non-uniform lengths of long reads. Therefore, mm2-plus delivers significant speedup over minimap2 across a range of alignment use cases. We benchmarked mm2-plus, mm2-fast [16], and minimap2 [21] for long-read mapping and all-vs-all read alignment tasks by using PacBio Hifi, Nanopore simplex, Nanopore duplex, and Nanopore ultra-long datasets from HG002 human genome (Supplementary Table S5). We used 48 threads of Cascade Lake CPU in this experiment. We demonstrate that mm2-fast and mm2-plus are faster than minimap2 across all use cases (Figure 7). The speedup of mm2-plus over minimap2 ranges from 1.32 *×* to 1.95 *×*. We occasionally see small improvements in mm2-plus over mm2-fast due to our addition of dynamic load balancing.

**Figure 7.**
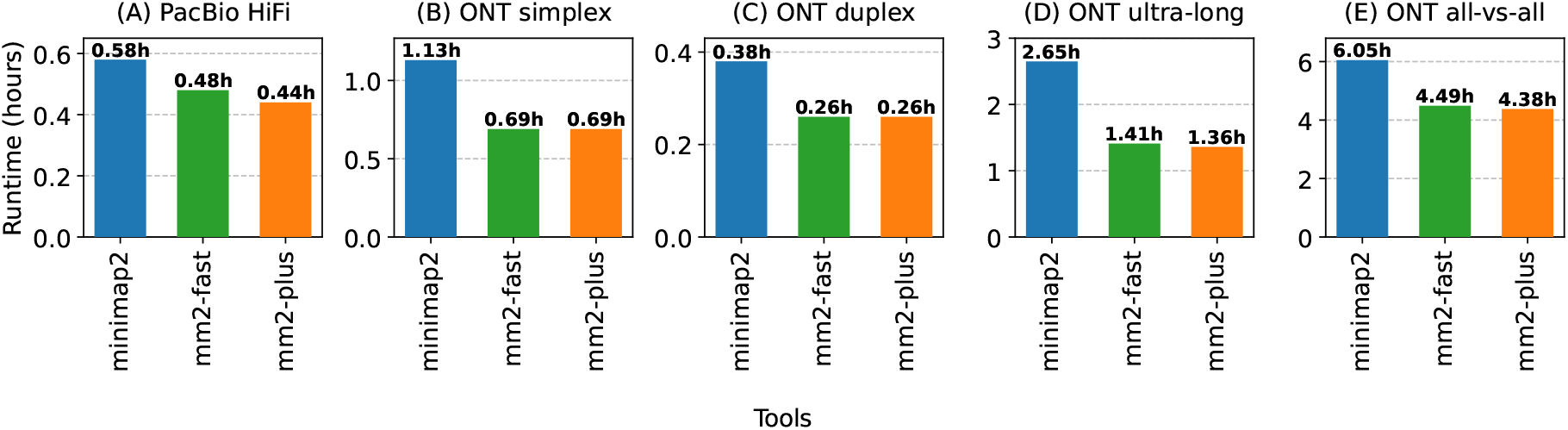
Comparison of runtimes of minimap2 [21], mm2-fast [16], and mm2-plus for long-read mapping and all-vs-all read alignment use-cases. We used whole-genome long-read sequencing data from Pacific Biosciences (PacBio) and Oxford Nanopore Technology (ONT) instruments. Panels (A)-(D) demonstrate speedup achieved while mapping long reads to CHM13 human genome [32]. Panel (E) demonstrates speedup achieved for all-vs-all alignment of ONT simplex reads. We ran each method using 48 threads on a Cascade Lake CPU. The commands used in this experiment are listed in Supplementary Table S3.

## 3 Discussion

Alignment of genome sequences is a routine computational task to study genome evolution across the tree of life. Within a species, alignments of fully phased genome assemblies enable discovery of the full spectrum of genetic variation [9, 26]. Improvements in per-base sequencing cost and accuracy are driving the increased adoption of long-read sequencing technologies in genomics. The number of genome assemblies to be analyzed is expected to increase proportionally. Furthermore, the assemblies generated using the latest long-read technologies comprise longer and fewer sequences, when compared to the assemblies generated from short-reads or noisy long reads [19]. In this work, we highlighted that the common approach for parallelizing genome alignment on multi-core processors is sub-optimal for aligning complete and near-complete assemblies. The parallelism available at the level of query sequences is limited, raising a need for an alternative approach.

We exclusively focused on improving one of the widely used genome aligners, minimap2 [21]. Our results revealed that the CPU utilization of minimap2 is consistently low across multiple datasets. To alleviate this, we developed mm2-plus by improving the various time consuming components of minimap2. We implemented several optimizations in mm2-plus that helped us achieve faster runtime without affecting the quality of alignments. These included faster algorithms for chaining, marking of primary chains, sorting and base-to-base alignment. Implementing these optimizations required a thorough understanding of the minimap2 source code, which was particularly challenging given that minimap2 consists of over 10,000 lines of C code. This paper also complements our earlier work on accelerating long-read mapping [16]. To make it convenient for users, we have ported the optimizations from [16] into mm2-plus. Therefore, mm2-plus can be utilized not only for whole-genome alignment but also for fast long-read mapping and all-vs-all read alignment. We have ensured that the output accuracy matches the accuracy of minimap2 (v2.28). Therefore, in any genomics workflow wherever alignment tasks are a bottleneck, users may benefit from using mm2-plus.

We expect our improvements to have a wider applicability beyond minimap2. Importantly, our optimizations can be adapted for any alignment method that employs seed-chain-extend heuristics. The proposed ideas could also be useful for accelerating pangenome graph construction because the graph construction requires progressive alignment of high-quality genome assemblies to intermediate pangenome graphs [14, 23].

## 4 Methods

### Faster parallel algorithm for chaining

The input to the chaining algorithm is a set of anchors, that is, short exact matches, between query sequences and the reference genome. The output of this step is a set of chains, where each chain comprises multiple anchors that can be combined to obtain a high-scoring alignment. In minimap2, sequences of the query genome are processed in batches. The query sequences in a batch are processed independently in parallel by utilizing one thread per sequence. The count of the query sequences in a batch depends on the batch size. The default batch size in minimap2 is 500 million bases. Processing a query genome in multiple batches rather than all the sequences at once helps in controlling the memory-usage.

Chaining of anchors is done using a dynamic programming algorithm, requiring *O*(*N* log *N*) time, where *N* denotes the number of anchors [1, 15]. Minimap2 uses a range-tree-based implementation for chaining [22]. The chaining step consumes a significant fraction of the overall time, as indicated by the profiling results (Figure 3). In mm2-plus, we ensured that the processing of each query sequence is parallelized across threads to achieve better CPU utilization and runtime. Note that each anchor is denoted by its (i) position on the query genome, (ii) position on the reference genome, and (iii) length of the match. Given the set of anchors from a query sequence, our parallel algorithm works by partitioning the anchors according to the reference genome sequence they match with. The number of partitions created equals the number of sequences in the reference genome. Subsequently, we perform independent chaining on each partition. This procedure does not affect the correctness because a chain cannot span over more than one reference sequence. We distribute the partitions across threads to perform chaining. We use dynamic assignment of the threads to each partition to handle non-uniform distribution of anchors across the reference sequences. We implemented this strategy to accelerate both the recursion and traceback phases of dynamic programming.

### Marking primary chains

The distinction between primary and secondary chains is required when a segment of a query genome aligns to two or more positions in the reference genome (Figure 1B). The output of the chaining algorithm is an array of anchor chains, denoted as 𝒞. Array 𝒞 is ordered in descending order of the chain scores. Suppose the number of chains in 𝒞 is *n*. We can identify the start and end coordinates of a chain over the query genome using the query start coordinate of the first anchor and the query end coordinate of the last anchor in the chain, respectively. For a chain *c*, let us denote *c*.*s* and *c*.*e* as its query start and query end coordinates, respectively. If there exist two or more chains having significant overlaps with each other, then the chain with the highest score is considered primary in minimap2, and the others are considered secondary. Before discussing our interval tree-based implementation, we review the quadratic-time algorithm used in minimap2.

For two chains *c*_*i*_ and *c*_*j*_, their *overlap fraction* is the ratio of their overlap length and the length of the shorter chain. Mathematically, we can write this as the ratio of max (0, min(*c*_*i*_.*e, c*_*j*_.*e*)− max(*c*_*i*_.*s, c*_*j*_.*s*)) to min(*c*_*i*_.*e* − *c*_*i*_.*s, c*_*j*_.*e*− *c*_*j*_.*s*). The algorithm in minimap2 works as following. Let us denote the set of primary chains as *Q*. Set *Q* is initially empty. We add the first chain from𝒞, i.e., the highest-scoring chain to *Q*. Subsequently, we consider the remaining chains in 𝒞 [2..*n*] sequentially. Chain𝒞 [*i*] is marked as secondary if there exists a chain in *Q* that has an overlap fraction above 0.5 with𝒞 [*i*]. Otherwise, 𝒞 [*i*] is marked as primary and added to set *Q*. Each iteration of the algorithm linearly scans set *Q*. Accordingly, this algorithm requires *O*(*n*^2^) time in the worse-case because |*Q* | can be as large as *n*.

In our implementation, we utilize an interval tree to avoid linear scans of *Q*. Interval tree is a standard data structure to store intervals. An interval tree comprising *m* intervals supports (i) addition of a new interval in *O*(log *m*) time, and (b) querying of all intervals that intersect with any given interval in *O*(log *m* + *k*) time, where *k* denotes the number of intervals produced by the query [7]. In mm2-plus, we use the same approach as minimap2, but maintain the intervals associated with the chains of set *Q* in an interval tree. While processing chain 𝒞 [*i*], we retrieve all the overlapping intervals in *Q* by making a query in the tree. As a result, each iteration now takes *O*(log | *Q* | + *k*) time. In our experiments, we found *k* ≪ *n*. Accordingly, this approach uses (*n* log *n*) time in practice.

### Accuracy validation

We evaluated the accuracy of genome alignments produced by mm2-plus using variant calling. For each dataset (Barley-Barley, Maize-Maize, Human-Human, Human-Bonobo), we used the alignment between the query genome and the reference genome to call variant using paftools (Commit: 0cc3cdc; https://github.com/lh3/minimap2/tree/master/misc). Paftools looks at the base-level differences in the sequence alignments and outputs substitution, insertion, and deletion variants. It considers both small and large variants. We assumed ground truth as the set of variants obtained from minimap2 whole-genome alignments. Subsequently, we measured the accuracy of variants derived from mm2-plus whole-genome alignments using bcftools (v1.20-58-gf6ac1c22; https://github.com/samtools/bcftools). The commands used for running paftools and bcftools are listed in Supplementary Table S3.

## Acknowledgements

This research is funded in part by the DBT/Wellcome Trust India Alliance Fellowship (grant number IA/I/23/2/506979), the Intel India Research Fellowship, and the National Supercomputing Mission India. We utilized computing resources available at the Indian Institute of Science, Intel Corporation, and the U.S. National Energy Research Scientific Computing Center.

Optimization Notice: Software and workloads used in performance tests may have been optimized for performance only on Intel microprocessors. Performance tests, such as SYSmark and MobileMark, are measured using specific computer systems, components, software, operations and functions. Any change to any of those factors may cause the results to vary. You should consult other information and performance tests to assist you in fully evaluating your contemplated purchases, including the performance of that product when combined with other products. For more information go to http://www.intel.com/performance. Intel, Xeon, and Intel Xeon Phi are trademarks of Intel Corporation in the U.S. and/or other countries.

## Supplementary Material

**Figure S1:**
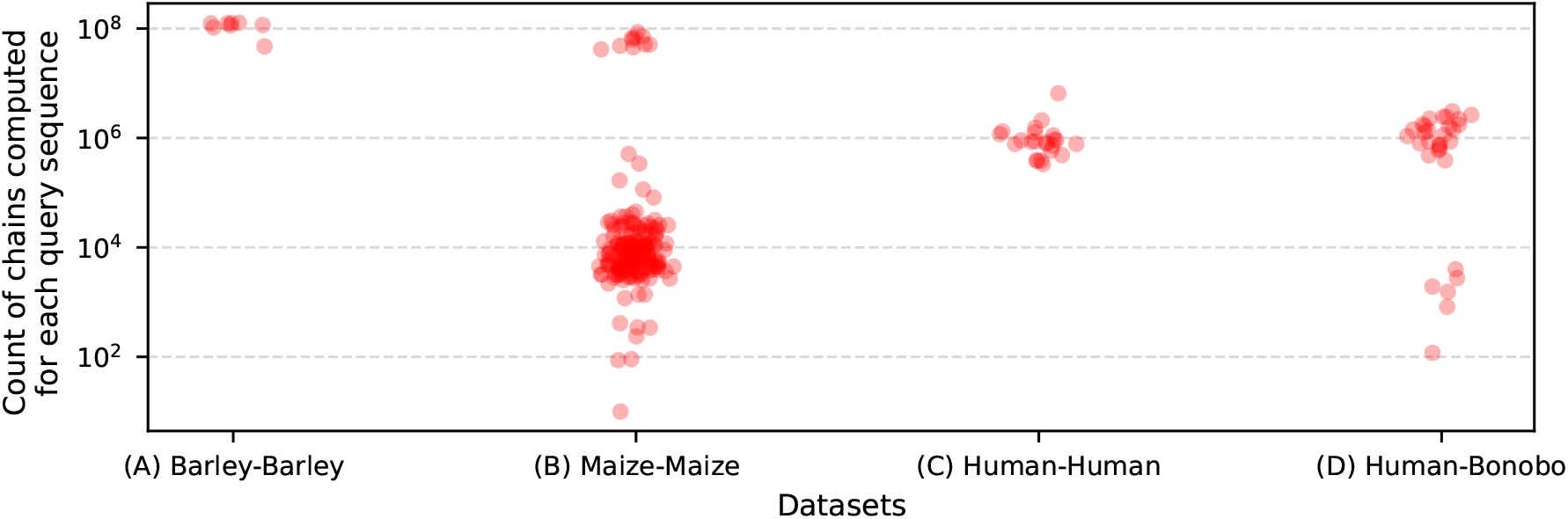
The plot illustrates the count of chains computed between each sequence of the query genome and the reference genome on various datasets.Each red circle represents a query sequence. These chains are further processed and classified as either primary or secondary in a subsequent step of the alignment workflow.

**Table S1:**
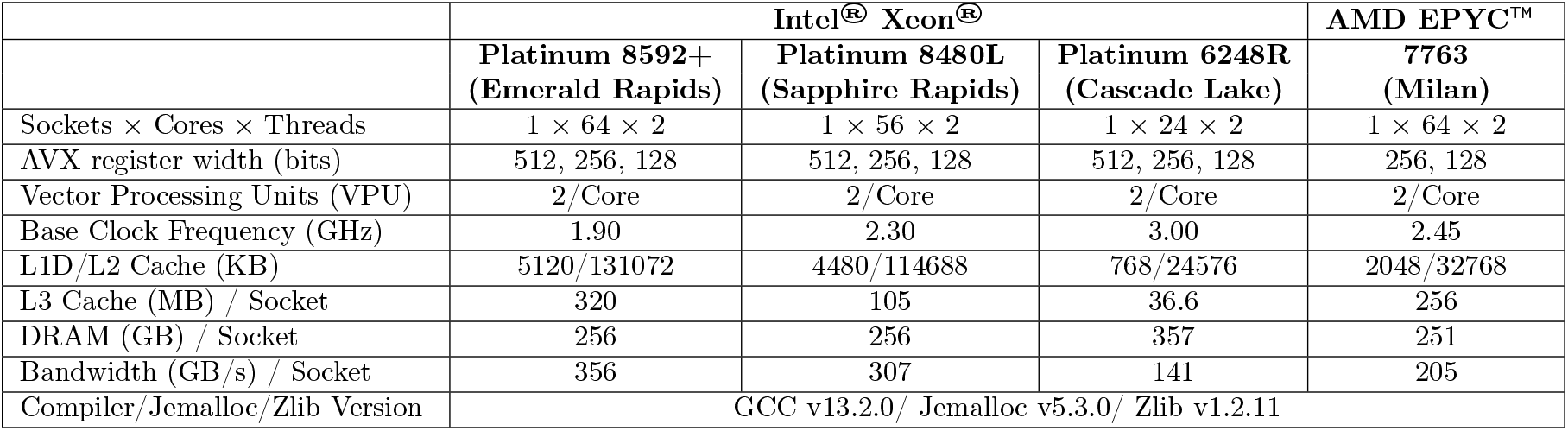
Overview of the architectural details for the four processors used in the experiments: Emerald Rapids, Sapphire Rapids, Cascade Lake, and Milan.

**Figure S2:**
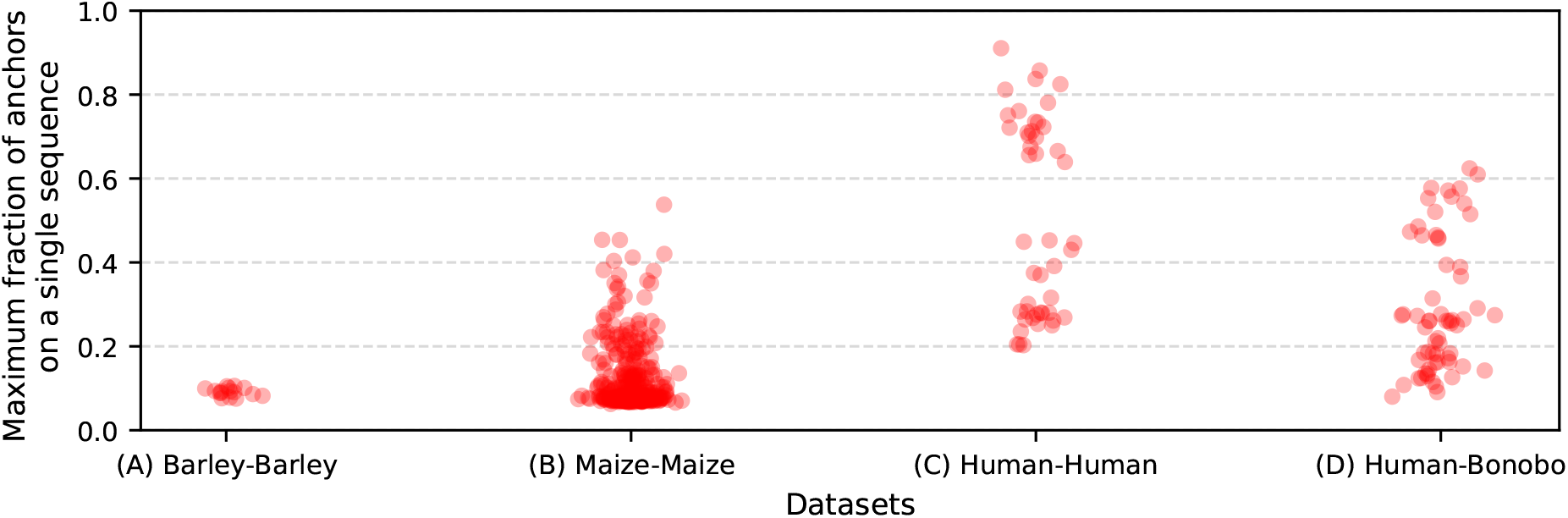
The output of the seeding stage is a set of anchors (exact matches) between each query sequence and the reference genome.The above plot illustrates the distribution of the count of anchors over various reference sequences. Specifically, for each query sequence (indicated as red circle), we measure the fraction of the maximum number of anchors on a reference sequence relative to the total number of anchors. A higher value of the fraction indicates that most anchors are located on a single reference sequence, that is, the distribution of anchors is skewed. A lower value indicates that the anchors are scattered across various reference sequences, that is, the distribution is more uniform. We observe low fraction values in plant genomes compared to animal genomes due to the higher presence of inter-chromosomal repeats and mobile elements.

**Figure S3:**
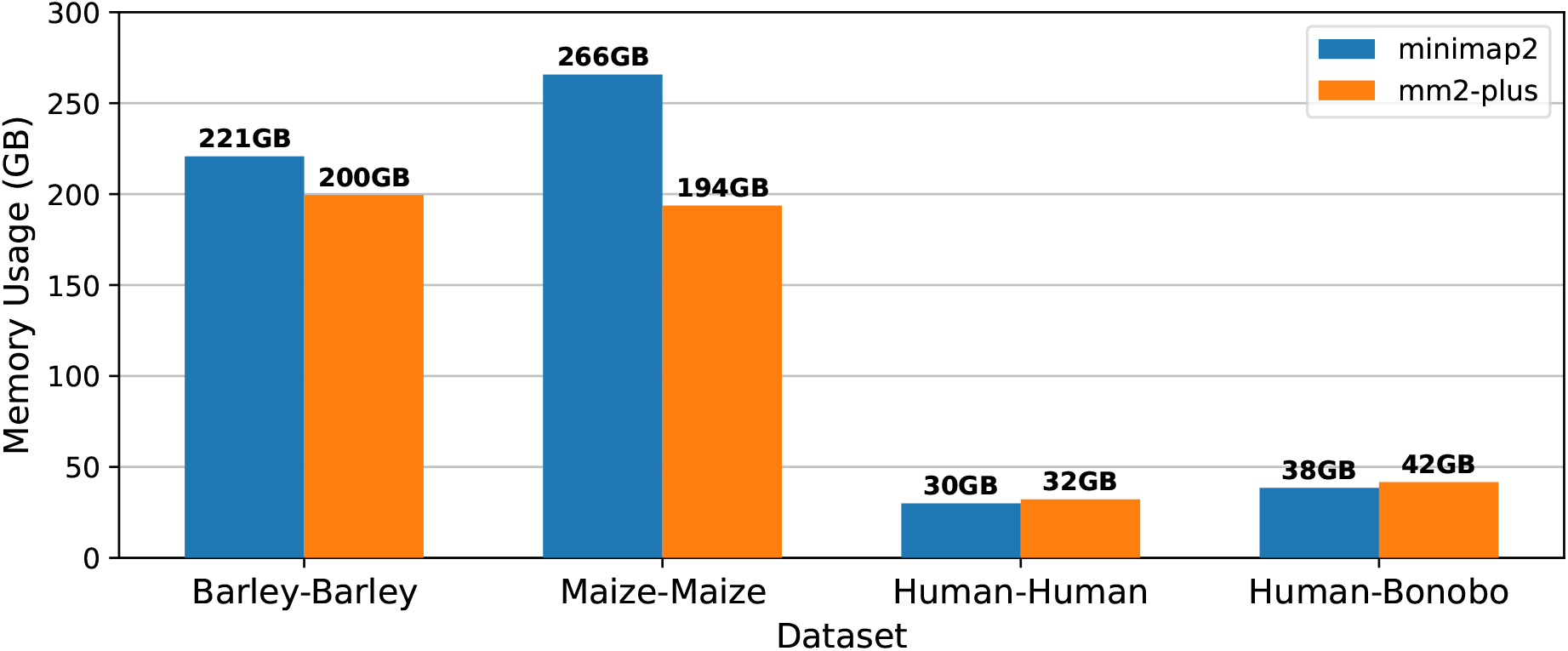
Memory usage comparison between minimap2 and mm2-plus. We conducted this experiment using 48 threads on a Cascade Lake CPU. All our optimizations were turned on in mm2-plus in this experiment.

**Table S2:** Description of the four datasets used in benchmark. In each dataset, we list the reference genome, the query genome and the NCBI accession numbers.

| Datasets | Reference |  | Query |  |
| --- | --- | --- | --- | --- |
|  | Genome | Accession Number | Genome | Accession Number |
| Human-Human | CHM13 v2.0 (Human) | GCA_009914755.4 | HG002 PATERNAL (Human) | GCA_018852605.2 |
| Human-Bonobo | CHM13 v2.0 (Human) | GCA_009914755.4 | mPanPan1 PATERNAL (Bonobo) | GCF_029289425.2 |
| Maize-Maize | Mo17 CAU 2.0 (Maize) | GCA_022117705.1 | W22 NRGENE 2.0 (Maize) | GCA_001644905.2 |
| Barley-Barley | Morex V3 (Barley) | GCA_904849725.1 | GPv1 (Barley) | GCA_902500625.1 |

**Table S3:**
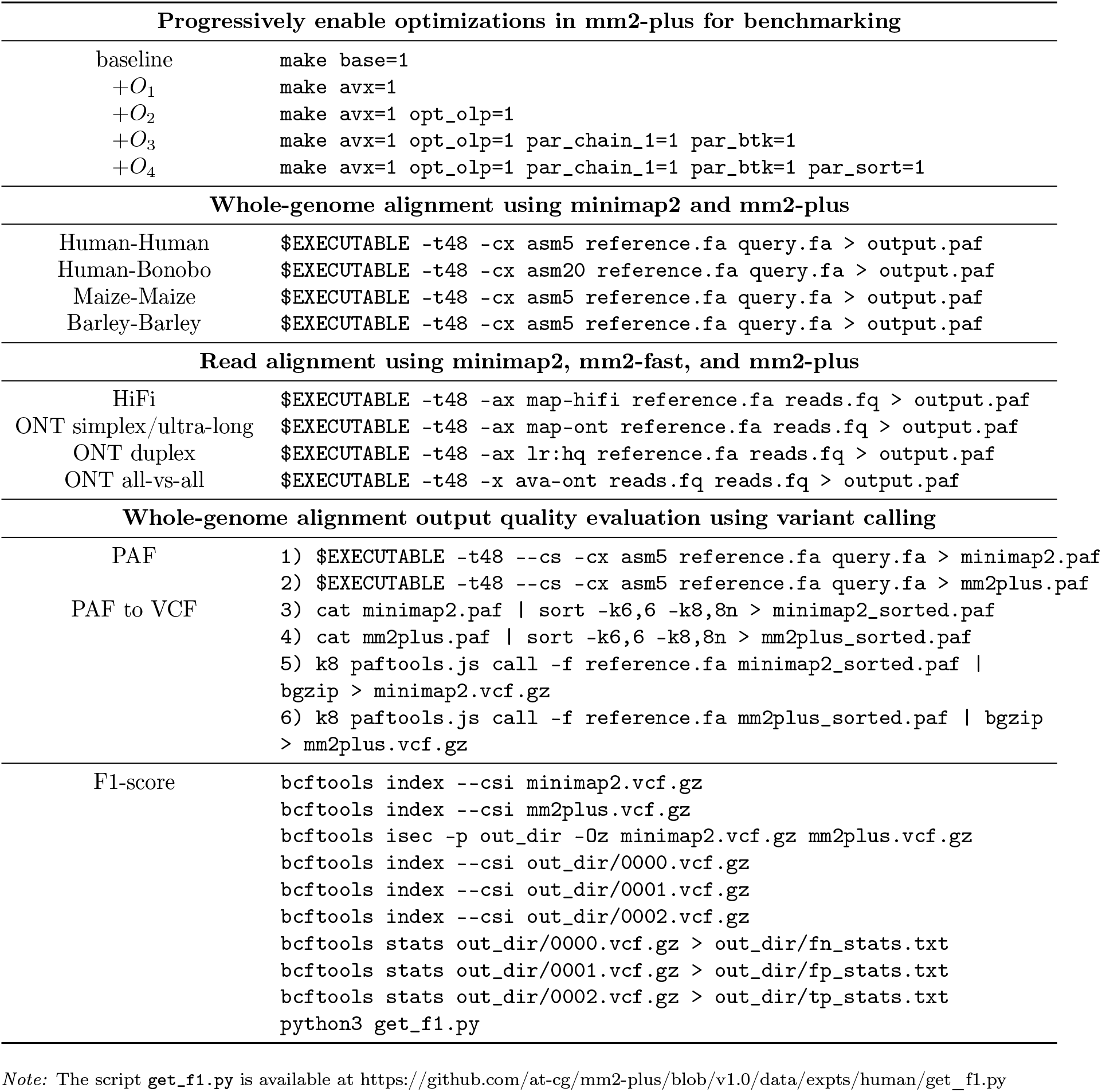
Commands used for running the experiments are listed in the table below. Additionally, we provide automated scripts to reproduce the results in the mm2-plus GitHub repository (https://github.com/at-cg/mm2-plus/tree/v1.0/data/expts).

**Table S4:** F1-scores comparing mm2-plus and minimap2 variant calls across four datasets. The high scores observed below indicate negligible differences in the alignment accuracy.

| Datasets | F1-score |
| --- | --- |
| Human-Human | 0.99990 |
| Human-Bonobo | 1.00000 |
| Maize-Maize | 1.00000 |
| Barley-Barley | 1.00000 |

**Table S5:**
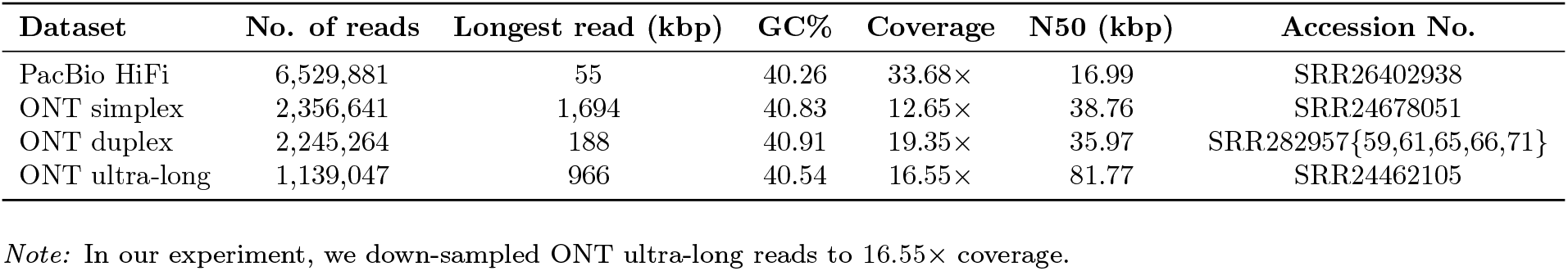
Summary statistics of long-read datasets. All read sets are sampled from HG002 human genome. For long-read mapping, we considered CHM13 T2T reference (GCA_009914755.4) as the reference genome.

